# Mitochondrial Biomarkers and Metabolic Syndrome in Bipolar Disorder

**DOI:** 10.1101/2023.12.13.571526

**Authors:** Kassandra A. Zachos, Jaehyoung Choi, Ophelia Godin, Timofei Chernega, Haejin Angela Kwak, Jae H. Jung, Bruno Aouizerate, Valérie Aubin, Frank Bellivier, Raoul Belzeaux R, Philippe Courtet, Caroline Dubertret, Bruno Etain, Emmanuel Haffen, Antoine Lefrere A, Pierre-Michel Llorca, Emilie Olié, Mircea Polosan, Ludovic Samalin, Raymund Schwan, Paul Roux, the FondaMental Academic Centers of Expertise in Bipolar Disorders (FACE-BD) Collaborators, Caroline Barau, Jean Romain Richard, Ryad Tamouza, Marion Leboyer, Ana C. Andreazza

## Abstract

**Importance:** Examining translatable mitochondrial blood-based biological markers to identify its association with metabolic diseases in bipolar disorder.

**Objective:** To test whether mitochondrial metabolites, mainly lactate, and cell-free circulating mitochondrial DNA are associated with markers of metabolic syndrome in bipolar disorder, hypothesizing higher lactate but unchanged cell-free circulating mitochondrial DNA levels in bipolar disorder patients with metabolic syndrome.

**Design:** In a cohort study, primary testing from the FondaMental Advanced Centers of Expertise for bipolar disorder was conducted, including baseline plasma samples and blinded observers for all experimentation and analysis.

**Setting:** The FondaMental Foundation coordinate a multicenter, multidisciplinary French networks aiming at creation of cohorts to improve identification of homogeneous subgroups of psychiatric disorders toward personalized treatments.

**Participants:** The FACE-BD primary testing cohort includes 837 stable bipolar disorder patients. The I-GIVE validation cohort consists of 235 participants: stable and acute bipolar patients, non-psychiatric controls, and acute schizophrenia patients. Participants were randomly selected based on biosample availability.

**Exposures:** All patients underwent the standard primary care within their center. No intentional exposures were part of this study.

**Main Outcome and Measures:** The primary outcome modelled an association with lactate and metabolic syndrome in this population. Reflective *a priori* hypothesis.

**Results:** Multivariable regression analyses show lactate association with triglycerides (Est= 0.072(0.023), p = 0.0065,), fasting glucose (Est = 12(0.025), p= 0.000015) and systolic (Est= 0.003(0.0013), p= 0.031) and diastolic blood pressure (Est = 0.0095±0.0017, p= 1.3e-7). Significantly higher levels of lactate were associated with presence of metabolic syndrome (Est = 0.17±0.049, p=0.00061) after adjusting for potential confounding factors. Mitochondrial-targeted metabolomics identified distinct metabolite profiles in patients with lactate presence and metabolic syndrome, differing from those without lactate changes but with metabolic syndrome. Circulating cell-free mitochondrial DNA was not associated with metabolic syndrome.

**Conclusion & Relevance:** This thorough analysis mitochondrial biomarkers indicate the associations with lactate and metabolic syndrome, whereas circulating cell-free mitochondrial DNA is limited in the context of metabolic syndrome. This study is relevant to improve the identification and stratification of bipolar patients with metabolic syndrome and provide potential personalized-therapeutic opportunities.

**Key Points:** *Question:* Can lactate, a mitochondrial metabolite, indicate metabolic syndrome in bipolar disorder?

*Finding:* In 837 stable bipolar disorder patients, we found high lactate levels significantly associated with metabolic syndrome, unlike circulating cell-free mitochondrial DNA. This pattern also appeared in acute bipolar and schizophrenia cases. Mitochondrial-targeted metabolomics distinguishes patients with high lactate and metabolic syndrome from those without lactate changes, but presence of metabolic syndrome.

*Meaning:* This research underscores lactate as a potential biomarker for identifying bipolar disorder patients with metabolic syndrome. It opens new avenues for personalized treatment strategies, leveraging mitochondrial metabolite profiling to improve patient stratification and therapeutic outcomes.

## Introduction

Clinical outcomes for bipolar disorder (BD) encompass a spectrum of comorbidities including high cardiovascular mortality^1^. This mood disorder is intimately associated with metabolic aberrations, such as high fasting glucose^2^, insulin resistance^2^, cholesterol levels^1^, leading to double rates of metabolic syndrome (MetS)^1, 3^. While several biological theories have been envisaged, compelling evidence of mitochondrial metabolic changes continue to grow. Henneman et al., as early as 1954, noted altered mitochondrial metabolism in psychotic patients, evidenced by increased blood lactate after glucose intake^4^. Kato and colleagues^5, 6^ studies reinforced this by identifying mitochondrial dysfunction in BD, observed through various markers in post-mortem brain tissues. Later studies confirmed these results, revealing gene expression changes in mitochondrial pathways and distinct brain imaging alterations. Studies also showed gene dysregulation in mitochondrial pathways for both BD and MDD^7, 8^ ^9–11^. Mitochondrial morphology were observed in BD post-mortem brain tissues, suggesting altered mitochondrial dynamics^12^, further confirmed by Choi et al. by using advanced data analysis leading to identify specific genes that categorize BD subgroups^13^, collectively underscoring mitochondrial dysfunction as a convergence pathway underpinning these metabolic clinical manifestations.

Dysfunctional mitochondria may arise from multifactorial origins (**Fig. 1**) leading to a metabolic shift, with consequent alterations of lipid biosynthesis and fatty acid oxidation^14–16^. This shift causes cells to be reliant on glycolysis for energy production, causing a build-up in lactate, which can act in competition with glucose as a fuel source and affect glucose uptake, which is implicated in the emergence of metabolic syndrome and may lead to the development of glucose intolerance^17, 18^. Concurrently, damaged mitochondrial structures result in release of mitochondrial DNA, known as circulating cell free mitochondrial DNA (ccf-mtDNA) to the periphery. Ccf-mtDNA is recognized as damage-associated molecular patterns (DAMPs).^19, 20^ This aberrant mitochondrial DNA outside the mitochondria triggers the TLR9 signaling cascade, amplifying/inducing NFKB-mediated pro-inflammatory gene expression and the activation of the NLRP3 inflammasome, culminating in chronic low-grade inflammation. MetS and inflammation are highly know to be trans-nosographically associated in patients with BD and other psychiatric diseases^21^.

**Figure 1.**
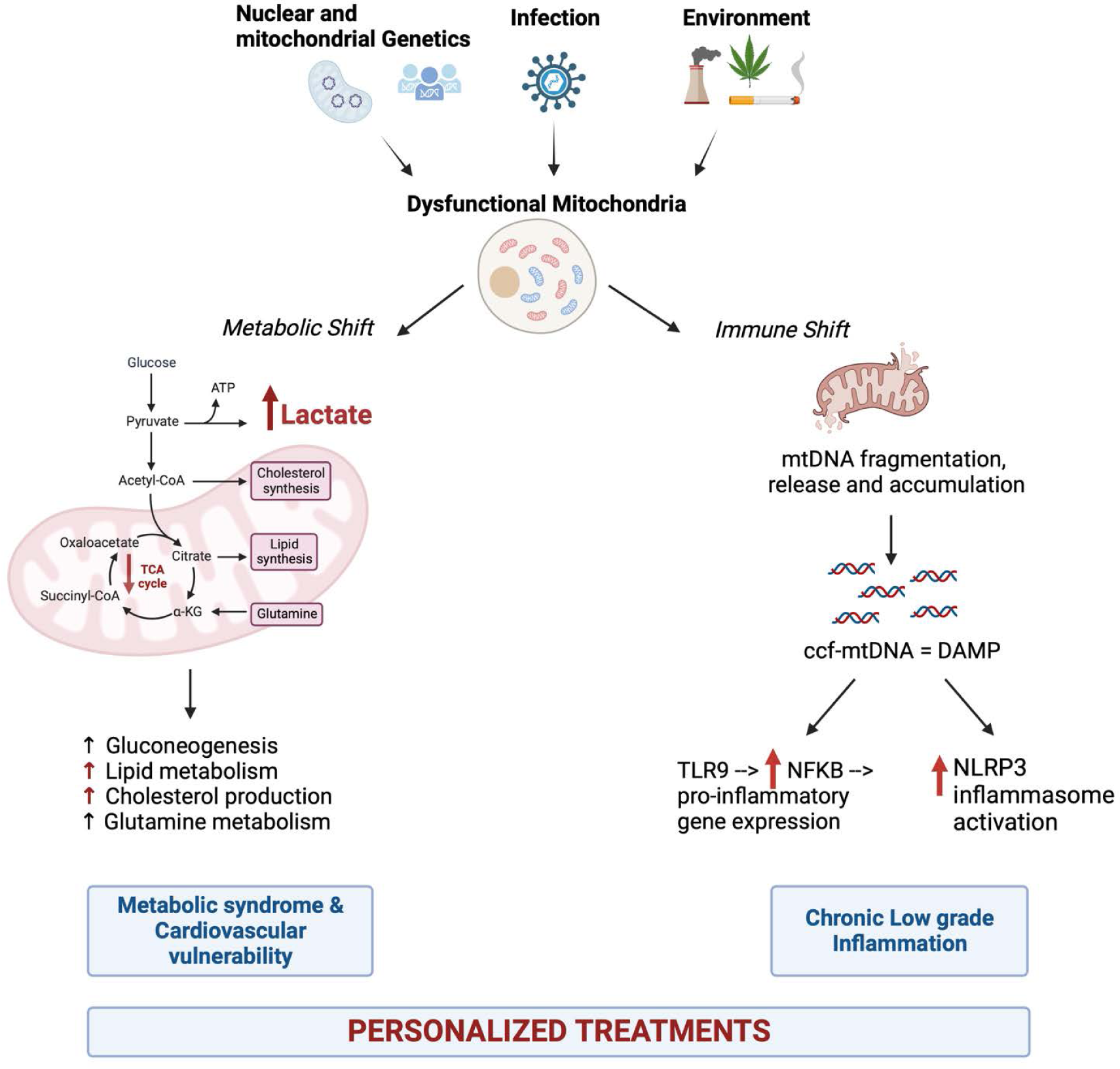
A model of mitochondrial dysfunction as a convergence pathway leading to metabolic or immune shift: towards personalized treatments.

Therefore, to further investigate the relationship between mitochondrial dysfunction and metabolic syndrome in BD, we examined key biological markers of mitochondrial dysfunction, lactate, along with mitochondrial targeted metabolomics, and ccf-mtDNA in patients with BD and determine the association of these markers with metabolic markers, clinical outcomes and MetS. Here, we included patients from the FondaMental Advanced Centers of Expertise in Bipolar Disorders (FACE-BD) cohort, a large and well-phenotype French cohort with stable BD patients and a smaller validation and replication cohort (I-GIVE) including stable and acute BD patients, as well as acute schizophrenia (SCZ) patients and non-psychiatric controls. We hypothesized that higher lactate and ccf-mtDNA levels will be associated with metabolic syndrome and c-reactive protein, as a marker of inflammation, respectively. Exploring biological markers that are easily translatable may ultimately demonstrate new ways for clinicians and researchers to potentially identify and treat metabolic changes, while offering new patient-specific therapeutic possibilities.

## Methods

### Study Population

The participant cohort comprised individuals undergoing evaluation at a collective of French healthcare facilities specializing in BD, a system instituted with the endorsement and financial aid of the French Ministry of Health and orchestrated by the FondaMental Foundation. Testing cohort (FACE-BD, **eFig. 1**) include stable outpatients diagnosed with BD(N=837) within the FondaMental Advanced Centers of Expertise (FACE-BD) cohort. Full description of population, characteristics and treatments can be found elsewhere^3, 23–26^. Validation cohort (N=255) consists of patients within stable or acute phases of BD (N=26 stable; N=75 acute) and during acute phase of SCZ (N=76) and non-psychiatric community controls (N=55) that were recruited as part of the French National granted I-GIVE (Immuno-Genetics, Inflammation, retro-Virus, Environment) cohort, for full description see^27^. Patients were evaluated, during hospitalization, for mania using the Young Mania Rating Scale (YMRS)^30^, Montgomery-Asberg Depression Rating Scale (MADRS) for depression^31^ and Positive and Negative Syndrome Scale (PANSS) was used to assess presence of psychotic symptoms^32^. IGIVE cohort subjects with acute phase had scores of MADRS above 17, YMRS above 8 or PANSS above 60^27^. **eFigure 1** and **eTables 1** and **2** shows the study design and clinical and demographics characteristics, respectively. After clinical phenotyping, biological samples were collected and processed by the biological research repository where samples were stored at -80°C until assayed.

### Metabolic Syndrome

MetS was defined according to the criteria of the International Diabetes Federation^28^, and requires the presence of ≥3 of the following criteria: high waist circumference (>90 cm for men and > 80 cm for women), hypertriglyceridemia (≥ 1.7 mmol/L or on lipid-lowering medication), low HDL cholesterol level (< 1.03 mmol/L in men and < 1.29 mmol/L in women), high blood pressure (≥ 130/85 mmHg or on antihypertensive medication), and high fasting glucose concentration (≥ 5.6 mmol/L or on glucose-lowering medication).

### Framingham Risk Score

The Framingham Risk Score is calculated using a point system, in which different scores are assigned to age, HDL-C, total cholesterol, blood pressure, diabetes and smoking status. This score is dependent on sex and current treatment and determines an individual’s 10-year CVD risk and heart age^29^.

### Circulating-cell free mitochondrial DNA (ccf-mtDNA)

was quantified in plasma using PCR amplification of ND1 and ND4, with methodological details described in **eMethods**. Experimenters were conducted blinded as well as raw analyses.

### Measurement of Lactate

Concentration of L-Lactate was measured using Cayman Chemical’s L-Lactate assay kit (Product number 700510) as per the manufacturer’s protocol for plasma samples and published elsewhere^22^. Experimenters were conducted blinded as well as raw analyses.

### Metabolomics

Plasma metabolomics profiling was conducted with collaborators at the University of Ottawa’s Metabolomics Core Facility. Targeted mitochondrial metabolomics were conducted of over 20 selected metabolites (**eTable3**), obtained from plasma were quantified by liquid chromatography mass spectrometry (LC-MS). Experimenters were blinded when conducting metabolomics and raw analyses.

### Statistical Analyses

Demographic and clinical variables were described in mean ± standard deviation or n (%). Fisher’s exact test and ANOVA was used to describe group-wise differences, where raw p-values are reported. The quantified ccf-mtDNA by copies of ND1 and ND4 showed linear correlation (Supplementary Methods Material), and natural-log transformation was applied to quantified ND1 (copies/μL) for statistical analyses. T-test, ANOVA, and Pearson’s correlation was used to statistically describe the relationship between lactate and ccf-mtDNA with demographic variables. Multivariable linear regression was used to describe the relationship between lactate, ccf-mtDNA and clinical variables, adjusting for age, sex, and BMI, where resulting p values were adjusted using Benjamini-Hochberg procedure. Pearson’s correlation test was used for the analysis of the correlation between metabolic markers. The threshold of p < 0.05 was used to determine statistical significance. Heatmap was visualized using seaborn (v0.12.2) in Python 3.8.5, where metabolites were z-transformed, then aggregated as group mean for visualization. Metabolomics hierarchical clustering was performed using Euclidian distance metric in Seaborn v0.12.2. Metabolome network was visualized in CytoScape, MetScape, and StringApp, and Pearson’s correlation matrix was used as the input for correlation mapping.

## Results

1. *Participant Demographics.* Of 837 patients in the testing cohort (FACE-BD), 556 (66.58%) are female, 46.66% (N=388) have a diagnosis of BD-type I. Among the FACE-BD cohort, the frequency of MetS was estimated to 17.76% (**eTable 1**). In the IGIVE cohort, demographical and clinical characteristics can be found in **eTable 2**.
2. *Levels of blood-based mitochondrial biomarkers in the FACE-BD cohort*. Regression analyses (**Fig. 2A,B**) adjusted for age, sex, and BMI, indicate that lactate is significantly associated with various markers related to MetS, including fasting glucose (**eFig. 2A**) triglycerides (**eFig. 2C**), and systolic and diastolic blood pressure (**Fig. 2A**). Significantly higher levels of lactate were also associated with frequency of MetS (**Fig. 2C**) whereas no significant association with clinical scales or CRP was observed.

Measures of ccf-mtDNA did not model the same associations with MetS. Ccf-mtDNA was not associated with MetS or with other metabolic markers (**Fig. 2A,B**). ccf-mtDNA was negatively correlated to one biological marker, specifically C-Reactive Protein (CRP) (**Fig. 2A**), a marker commonly utilized clinically to represent acute phase response and low-grade inflammation in many different physiological and psychiatric diseases^33^. ccf-mtDNA levels was positively associated (**Fig. 2B**) with manic symptoms (YMRS) while lactate levels did not demonstrate this, suggesting a potential association of lactate with metabolism and ccf-mtDNA with symptomatology, however further investigation is warranted.

**Figure 2.**
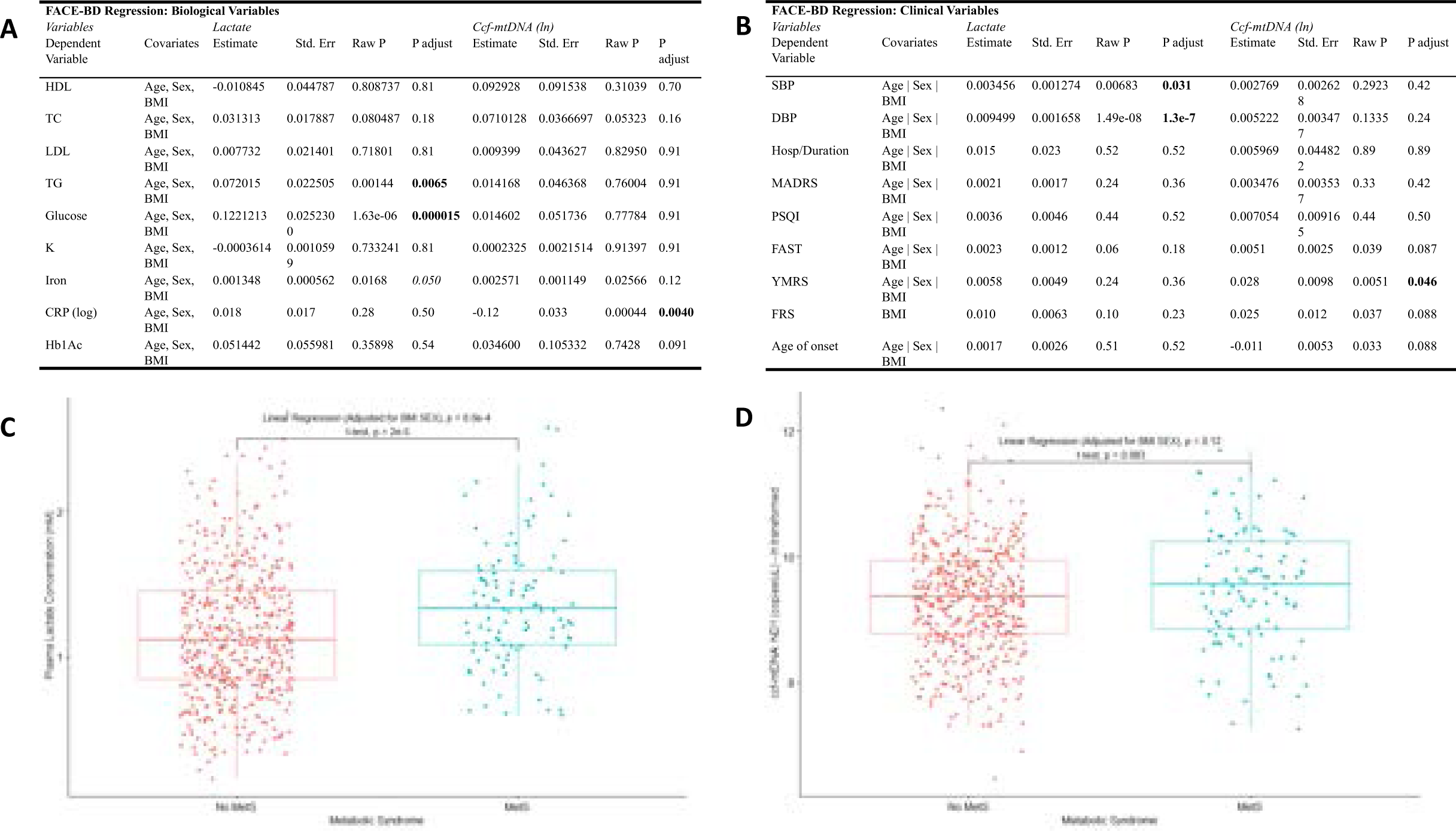
Regression analysis model for association of mitochondrial-blood based biomarkers with biological (A) and clinical (B) variable in the FACE-BD cohort. Association of peripheral levels of lactate (C) and circulating cell free mitochondrial DNA (ccf-mtDNA, D) with metabolic syndrome (MetS) in stable patients with bipolar disorder. High-density lipoprotein (HDL), total cholesterol (TC), triglycerides (TG), potassium (K), c-reactive protein (CRP), hemoglobin A1C (HbA1C), systolic blood pressure (SBP), diastolic blood pressure (DBP), hospitalization (Hosp); Young Mania Rating scale (YMRS); Functional Assessment Short Test (FAST); Farmington risk score (FRS).

Finally, we utilized relevant data to calculate the Framingham Risk score, a predictive tool developed for coronary heart disease, for individuals of the FACE-BD cohort ^29^. While there is nominal elevation of ccf-mtDNA copies with higher Framingham Risk Score, no significant association where observed (**Fig. 2B**).

### 2.1 Mitochondrial-Targeted Metabolomics in FACE-BD

In efforts to further elucidate the metabolic relationship present between lactate and MetS, we conducted mitochondrial targeted metabolomics to examine, several relevant metabolites involved in mitochondrial metabolism in stable BD patients. In total, this subgroup analysis included 100 patients, 50 with clinically defined low levels of lactate (<1mmol/L) and 50 patients with high lactate levels (>1.8mmol/L) (**Fig. 3A**). Just over 20 metabolites were selected for examination, but ultimately 16 metabolites were included in our final analyses due to a lack of signal (**eTable 3**). The frequency of MetS was almost 2 times higher in the high lactate group compared to the low lactate group although not significantly associated (x^2^=3.05; p=0.08) (**Fig. 3A**). While stratifying by metabolic syndrome or lactate levels alone yielded no significant mitochondrial associations (**Fig. 3B**), combining both factors pinpointed specific mitochondrial metabolites (**Fig. 3C**). Metabolite profiles between the high and low lactate groups display stronger than the groups with the presence of MetS. Notably, patients with elevated lactate and metabolic syndrome had higher citrate and alpha-ketoglutarate (AKG) levels than those with lower lactate (**Fig. 3C**). A metabolite correlation analysis emphasizes the association of AKG in this pathway (**Fig. 3E**). Conducting metabolomics emphasizes the relationship of lactate and MetS, while simultaneously provides insight on potential underlying mechanisms within individuals with BD and MetS. Mitochondrial metabolites showed a general trend towards inverse relationships with YMRS (**eFig. 3**).

3. *Replication Analysis: Mitochondrial Biomarkers in FACE-BD & I-GIVE*. To strengthen our analysis, we conducted a replication analysis utilizing age and sex-matched stable BD patients from a separate cohort, the I-GIVE cohort. Throughout the study, our inter- and intraplate variability remains relatively low for both lactate and ccf-mtDNA (**Fig. 4A**). Then we selected age and sex matched stable BD patients from both cohorts, it is evident that both plasma lactate (**Fig. 4B**) and ccf-mtDNA (**Fig. 4C**) levels remain consistent throughout both BD populations. Finally, we compare among the same individuals (N=100) lactate levels measured via mass spectroscopy and spectrophotometry showing a strong positive correlation (**Fig. 4D**).
4. *Validation Analysis: Blood-based mitochondrial biomarkers in I-GIVE*. Further we investigated the levels of lactate and ccf-mtDNA in patients with BD or SCZ during acute phase of the illness (**eTable 4**). Intriguingly, acute BD patients of showed significantly elevated lactate levels in comparison to stable BD patients and non-psychiatric controls (**Fig. 5A**). Notably, when compared to non-psychiatric controls lactate levels were also elevated in stable BD patients (**Fig. 5A**). Acute SCZ patients showed similar levels of lactate when compared to acute BD patients (**Fig. 5A**). We explored the relationship of lactate levels and clinical presentation of depression, hypomania or mania and observed no relationship, however the study was not designed to answer this question (**eTable 5**). No differences were observed for ccf-mtDNA levels in patients compared to non-psychiatric controls. Finally, we evaluate, in the IGive cohort, whether mitochondrial DNA copy number (previously published in these patients^27^) could affect the release of ccf-mtDNA, by creating a mt copy number to ccf-mtDNA ratio, no significant differences were observed across participants groups (**Fig. 5E**). To further explore sensitivity and specificity of lactate, we performed a receiver operating characteristic curve (ROC) showing that lactate differentiated between patients with acute psychiatric disorder compared to non-psychiatric controls (**Fig. 5F-H, eFig. 4**).

**Figure 3.**
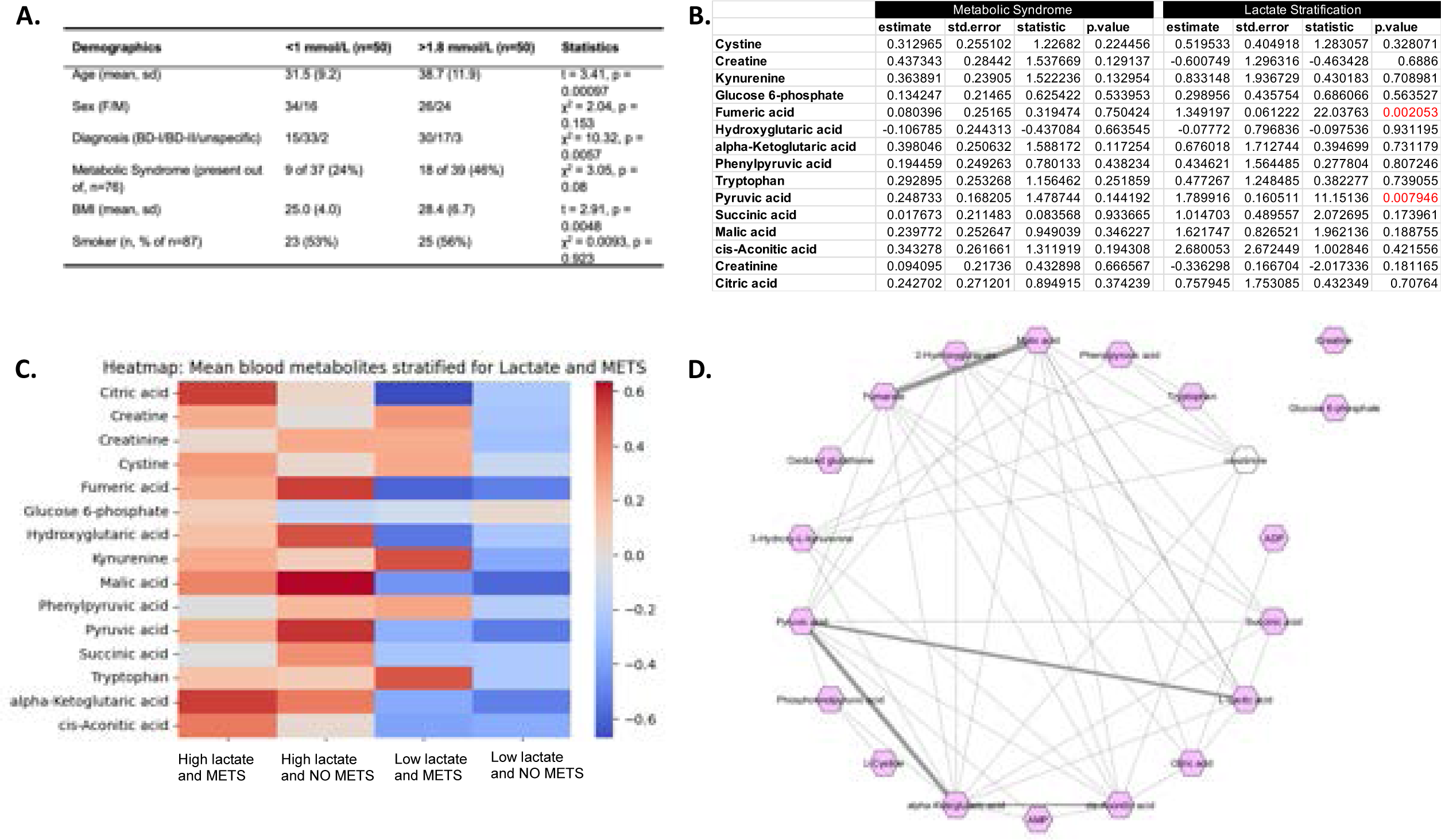
Mitochondrial targeted metabolomics in patients with BD (FACE-BD). A. Demographic distribution of patients with low (<1 mmol/L) and high (> 1.8 mmol/L) levels of lactate. B. Statistical analysis of mitochondrial metabolites by stratification via metabolic syndrome or lactate levels. C. Heatmap of mitochondrial blood metabolites stratified by lactate and metabolic syndrome (METS). D. Metabolites correlation analysis.

**Figure 4.**
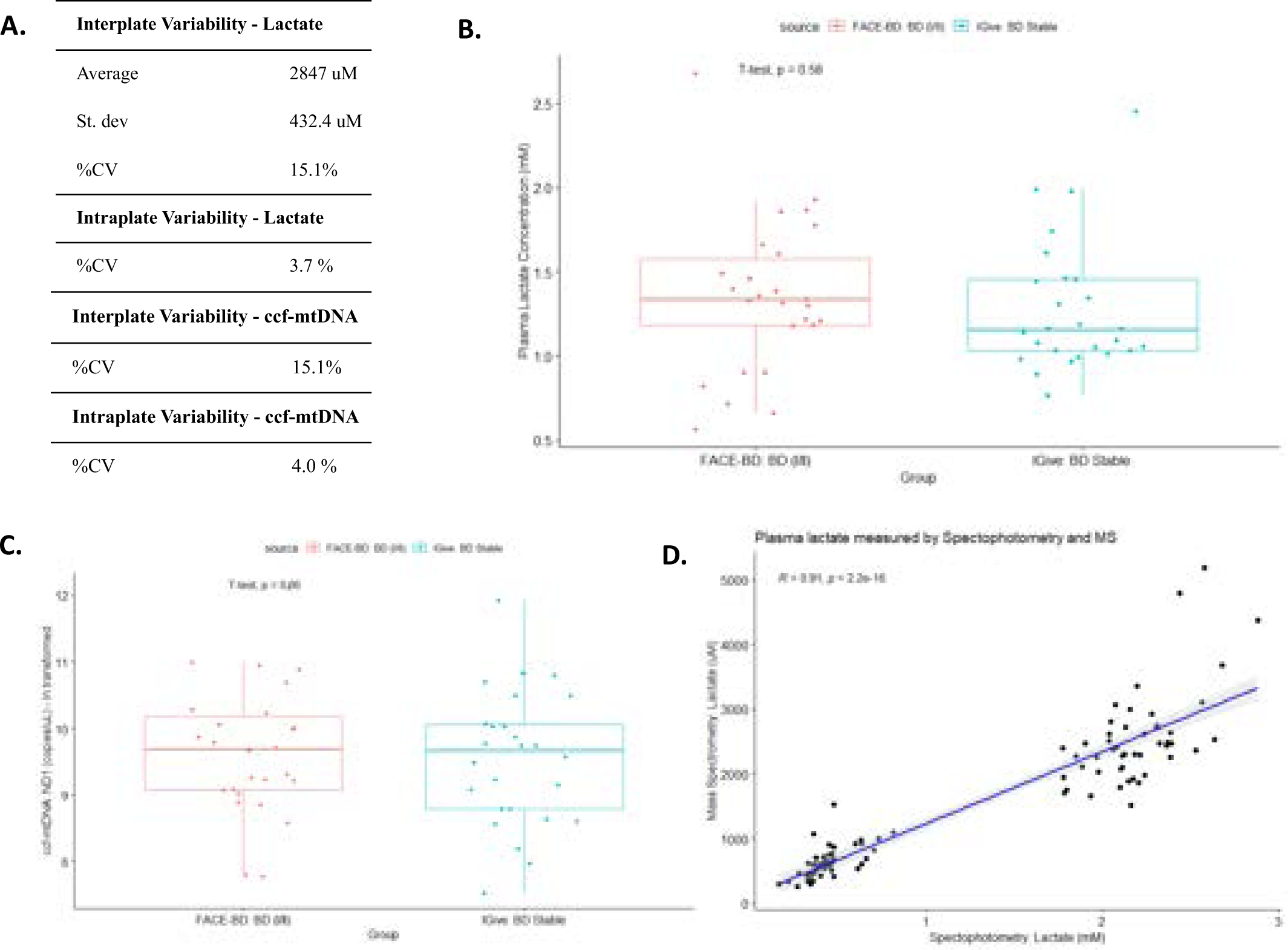
Age-sex matched replication analysis of mitochondrial-blood based biomarkers. A. Interplate and intraplate variability of lactate and circulating cell free mitochondrial DNA (ccf-mtDNA). B. Levels of lactate across FACE-BD and iGIVE cohort age-sex matched stable bipolar disorder (BD) patients. C. Levels of ccf-mtDNA across FACE-BD and iGIVE cohort age-sex matched stable bipolar disorder (BD) patients. D. Correlation of lacate levels measured by spectrophotometric measurement and liquid chromatography mass spectrometry (LC-MS).

**Figure 5.**
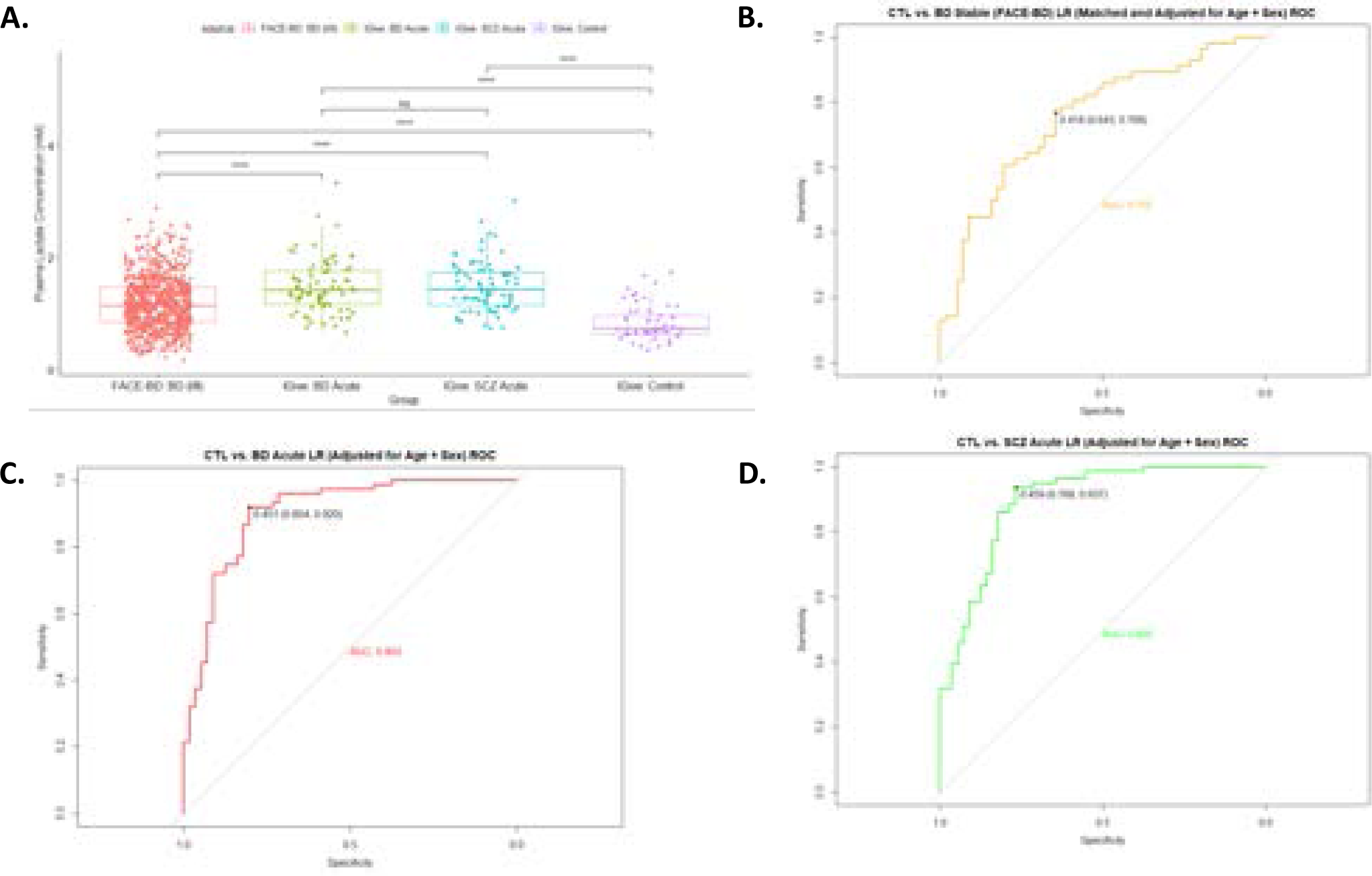
Levels of lactate (A) across acute patients with bipolar disorder (BD) and schizophrenia (SCZ) and non-psychiatric controls (controls) and (B - D) sensitivity and specificity analysis.

## Discussion

In the first study of its kind, utilizing a large cohort of stable BD patients, we were able to describe the utility of lactate as a metabolism in BD and the potential of mitochondrial metabolites to identify a distinct signature between patients with MetS and elevated levels of lactate compared to those with MetS only. Furthermore, lactate levels remain consistent among stable BD patients in both the FACE-BD and I-GIVE cohorts but are notably significantly lower in non-psychiatric controls and higher in both acute BD patients and acute SCZ patients (I-GIVE cohort). While identifying that ccf-mtDNA is not a marker of metabolism shift in patients with BD (**Fig. 1**). Our data collectively demonstrate the importance of mitochondrial metabolites in identifying homogenous patient groups likely to benefit from targeted mitochondrial metabolic interventions.

Lactate is a well-known product of glycolysis, that is critical in the homeostasis of metabolism^34, 35^. However, in a state of mitochondrial dysfunction and anaerobic conditions, there is a decrease in efficiency of the Krebs cycle and electron transport chain, forcing glycolysis to compensate to produce sufficient ATP for the organism to function^18^. In this case, pyruvate is reduced by NADH into lactate in high amounts and intracellular and extracellular lactate accumulation occurs, and this relationship is emphasized in the correlation analysis conducted^36^. Glucose fermentation into lactate is reported to occur at a faster rate than glucose oxidation by the mitochondria, linking this rapid increase in lactate levels as marker of circulatory dysfunction, tissue damage and metastasis in cancer^37, 38^. Lactic acidosis that occurs from lactate accumulation simultaneously promotes metastasis while inhibiting antitumor immune responses^39^. In psychiatric diseases, including mood disorders, autism and schizophrenia, elevated lactate levels have consistently been demonstrated in serum, brain and cerebrospinal fluid^40, 41^. Individuals with T2D plasma lactate levels were raised compared to non-disease controls, and associated with increased fasting glucose levels, lower HDL cholesterol levels and triglycerides^18, 37^.

Excess citrate, transported from the mitochondrial matrix to the cytosol, can suppress aerobic glycolysis, TCA cycle, and fatty acid breakdown, influencing metabolic diseases^42^. Moreover, heightened AKG levels are known to boost fatty acid production and elevate stored triglycerides^43^. AKG acts as a rate limiting step in the TCA cycle as glutamine rewiring can occur to convert AKG back into citrate in hypoxic cells or in the event of dysfunctional mitochondria, which may explain this link exhibited in BD patients with high lactate and metabolic syndrome^44–46^. In individuals with low lactate with metabolic syndrome, kynurenine and tryptophan were relatively elevated. Tryptophan is an essential amino acid metabolized by the kynurenine pathway, which can lead to the formation of quinolinic acid and nicotinamide adenine dinucleotide (NAD)^47^. The first of two rate limiting enzymes in this pathway, indoleamine-2,3-dioxygenase, has been theorized to play a role in major depressive disorder. The second enzyme, tryptophan 2,3-dioxygenase, is inhibited by glucose intake in the liver and downstream enzymes have been positively correlated with impaired glucose tolerance^47^.

Circulating-cell free mitochondrial DNA (ccf-mtDNA) occurs when dysfunctional mitochondria fragment and release this fragmented DNA into the periphery^48^. Increased ROS production can trigger released of its DNA into the cytosol through the mitochondrial transition channel opening^48^. Cell death occurs, typically via apoptosis, and the mitochondrial DNA accumulates in the periphery and acts as a DAMP triggering the TLR9 signaling cascade and formation of the NLRP3 inflammasome. NLRP3 activation is known to play a role in immune and autoimmune disease, both type 1 and 2 diabetes, cancer, and central nervous system diseases, including BD and SCZ^49^. While there was an observed link with ccf-mtDNA and CRP, further exploration in an immune-focused model, incorporating more markers of inflammation, is warranted.

These underlying mechanisms discussed are highly variable and speak to the clinical heterogeneity demonstrated in individuals with bipolar disorder. To best understand the etiology of the disease, and improve therapeutic outcomes, precision medicine requires thoughtful consideration. Tryptophan and lactate have been closely linked to the gut microbiome, altering their overall metabolism and plasma levels, providing a wide array of therapeutic targets to be explored^47, 50^. The ketogenic diet has been examined in the context of insulin resistance in BD patients as it provides ketones as an alternative mitochondrial fuel source^51^. Finally, the use of AKG as a nutritional supplement has been more commonly investigated to improve ATP supply, but this may only be useful in BD patients without metabolic syndrome, or those with lower lactate levels^45, 52^.

Within every study there are limitations, thus results should be interpreted with caution through lenses of investigational science. It is important to note that our primary cohort was sufficient in sample size, however our replication and validation cohort were comparatively much smaller, warranting further studies on additional cohorts to strengthen this model. All patients’ samples were collected after fasting; however, we had not accounted for lifestyle and diet. Plasma lactate was blindly collected, measured, and analyzed; however, the lactate measurements were not measured with fresh plasma. Rather, plasma was stored immediately at -80° Celsius and securely preserved until ready for use. Experiments were conducted in batches and samples were properly thawed on ice, to ensure quality of samples were maintained throughout. Metabolomic samples underwent a single freeze-thaw cycle, however all samples underwent the same cycle and thus samples were all handled comparatively. To ensure reliability of the experiments we compared levels of mitochondrial biomarkers across cohorts and compared levels of lactate using two different technologies (i.e., mass spectrometry and spectrophotometry) demonstrating accuracy of lactate levels. We performed ROC analysis to explore the sensitivity and specificity of lactate, caution should be applied in absence of additional testing sets the presented data is a statistical description of the cohorts we have examined.

In conclusion, our study marks a significant advancement in bipolar disorder research by revealing the critical role of lactate and mitochondrial metabolites in distinguishing metabolic profiles among patients. Demonstrating consistent lactate levels across BD cohorts and its differential levels among acute versus stable patients and non-psychiatric controls, we highlight its potential as a metabolic surrogate biomarker. Our findings also clarify that circulating cell-free mitochondrial DNA (ccf-mtDNA) does not mark metabolic shifts in BD, adding a new dimension to our understanding of mitochondrial dynamics in psychiatric conditions. These insights pave the way for personalized treatment strategies in bipolar disorder, emphasizing the importance of mitochondrial dynamics in psychiatric and metabolic health. This research opens new avenues for targeted interventions, promising a future of precision medicine tailored to individual metabolic and psychiatric needs^49^.

## Acknowledgements

We thank the support of the Baszucki Brain Fund for the financial support towards this study and the Mitochondrial Innovation Initiative (MITO2i) for support KZ scholarship. Agence Nationale de la Recherche (ANR-11-IDEX-0004-02 and ANR-10-COHO-10-01and ERANET Neuron-ANR-18-0008-01), INSERM (Institut National de la Santé et de la Recherche Médicale) and Fondation FondaMental (www.fondation-fondamental.org). We thank all the patients and healthy controls who participated to this study, the clinical teams from the University department of psychiatry and addictology of Henri Mondor hospital (DMU IMPACT and FHU ADAPT) as well as the team from the Biobank and CIC of Henri Mondor Hospital (AP-HP). Metabolites were analysed at the University of Ottawa Metabolomics Core Facility. This facility is supported by the Terry Fox Foundation and Ottawa University.

## Online-Only Supplemental Material

### Supplementary Tables

**eTable 1.**
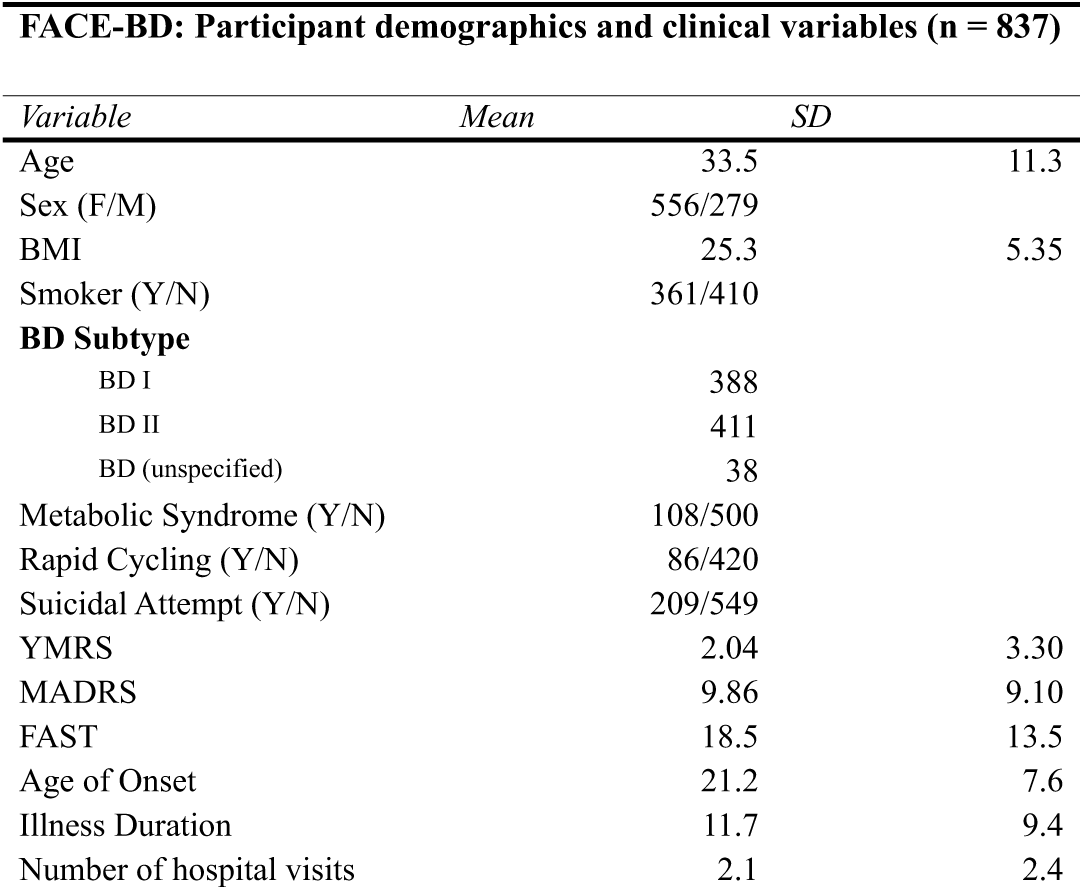
FACE-BD cohort clinical and demographics characteristics.

**eTable 2.**
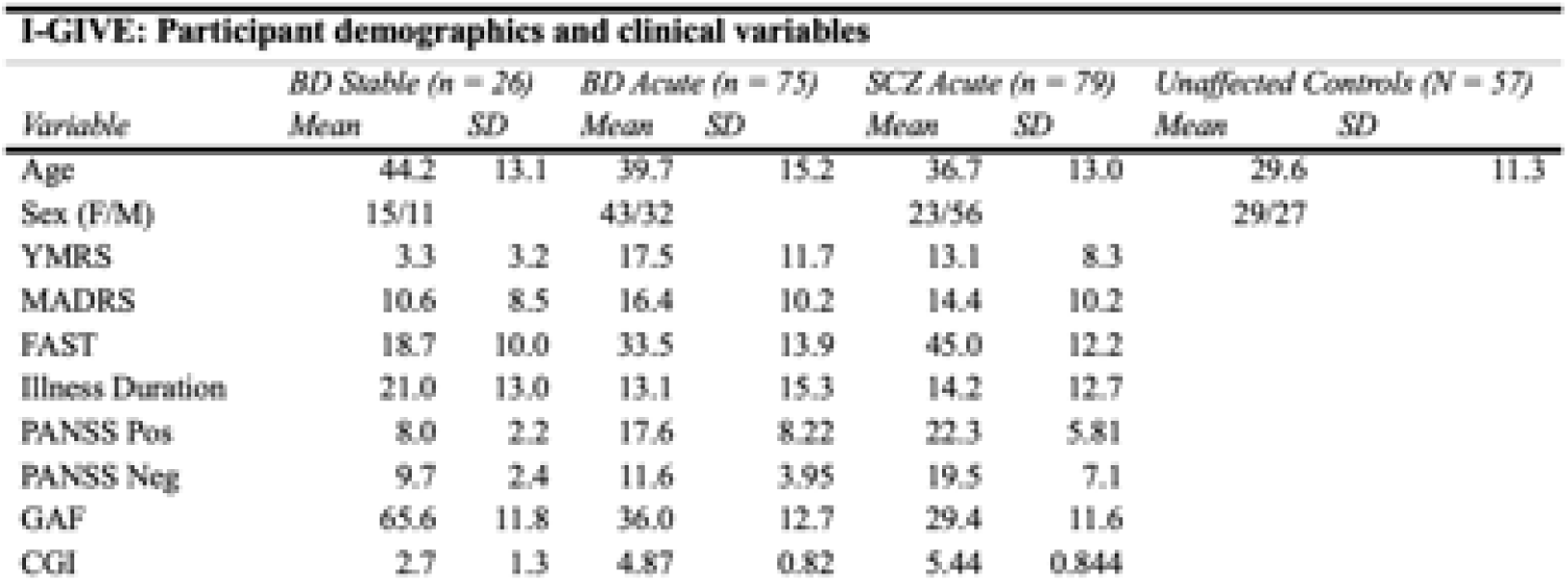
iGIVE cohort clinical and demographics characteristics.

**eTable 3.**
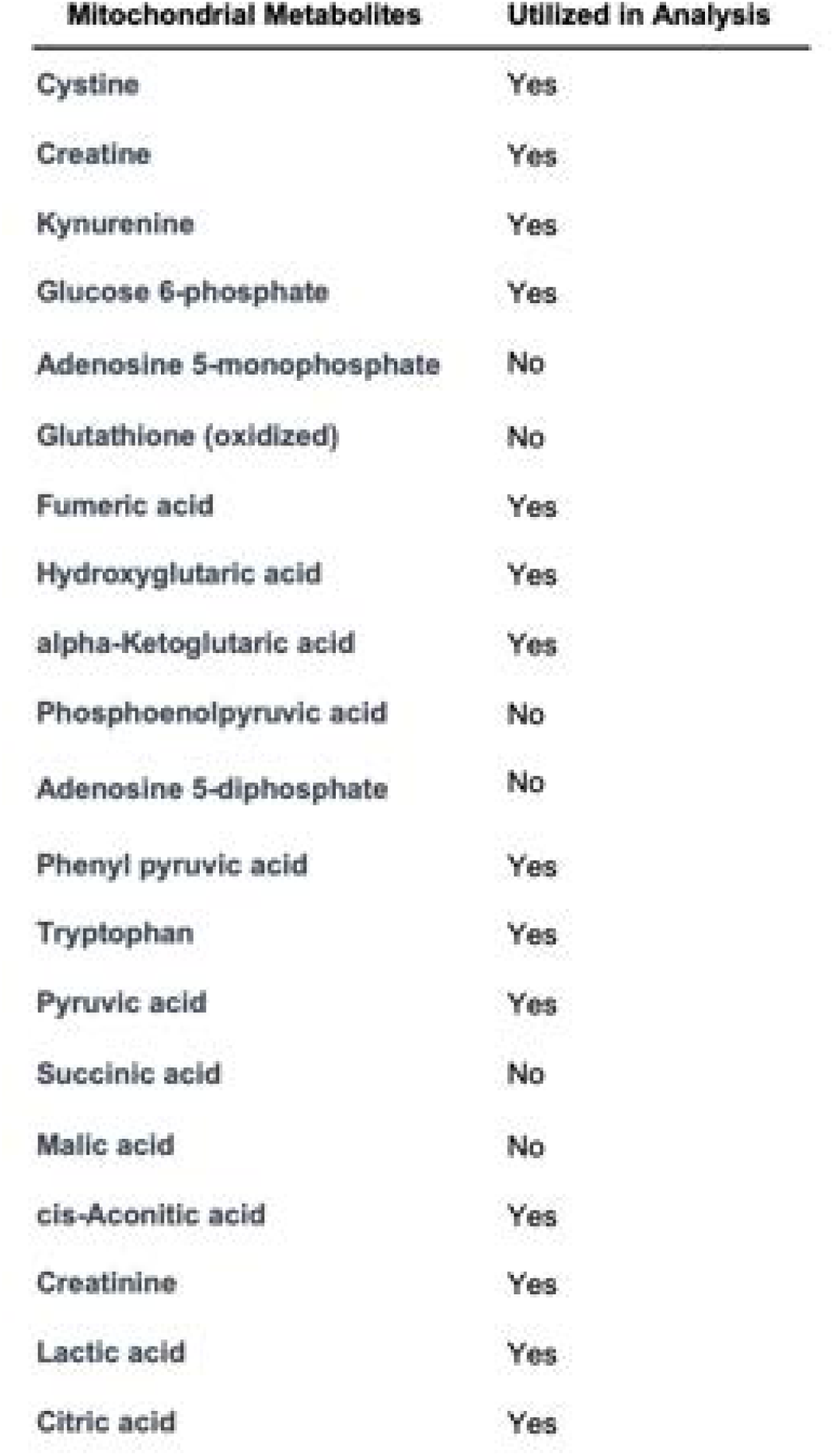
Mitochondrial metabolites assessed via liquid chromatography mass spectrometry (LC-MS), metabolites with lower limit of detection were not included in the final statistical modeling.

**Mitochondrial metabolites assessed via liquid chromatography mass spectrometry (LC-MS).**
| **Mitochondrial Metabolites** | **Utilized in Analysis** |
| --- | --- |
| Cystine | Yes |
| Creatine | Yes |
| Kynurenine | Yes |
| Glucose 6-phosphate | Yes |
| Adenosine 5-monophosphate | No |
| Glutathione (oxidized) | No |
| Fumaric acid | Yes |
| Hydroxyglutaric acid | Yes |
| alpha-Ketoglutaric acid | Yes |
| Phosphoenolpyruvic acid | No |
| Adenosine 5-diphosphate | No |
| Phenyl pyruvic acid | Yes |
| Tryptophan | Yes |
| Pyruvic acid | Yes |
| Succinic acid | No |
| Malic acid | No |
| cis-Aconitic acid | Yes |
| Creatinine | Yes |
| Lactic acid | Yes |
| Citric acid | Yes |

**eTable 4.**
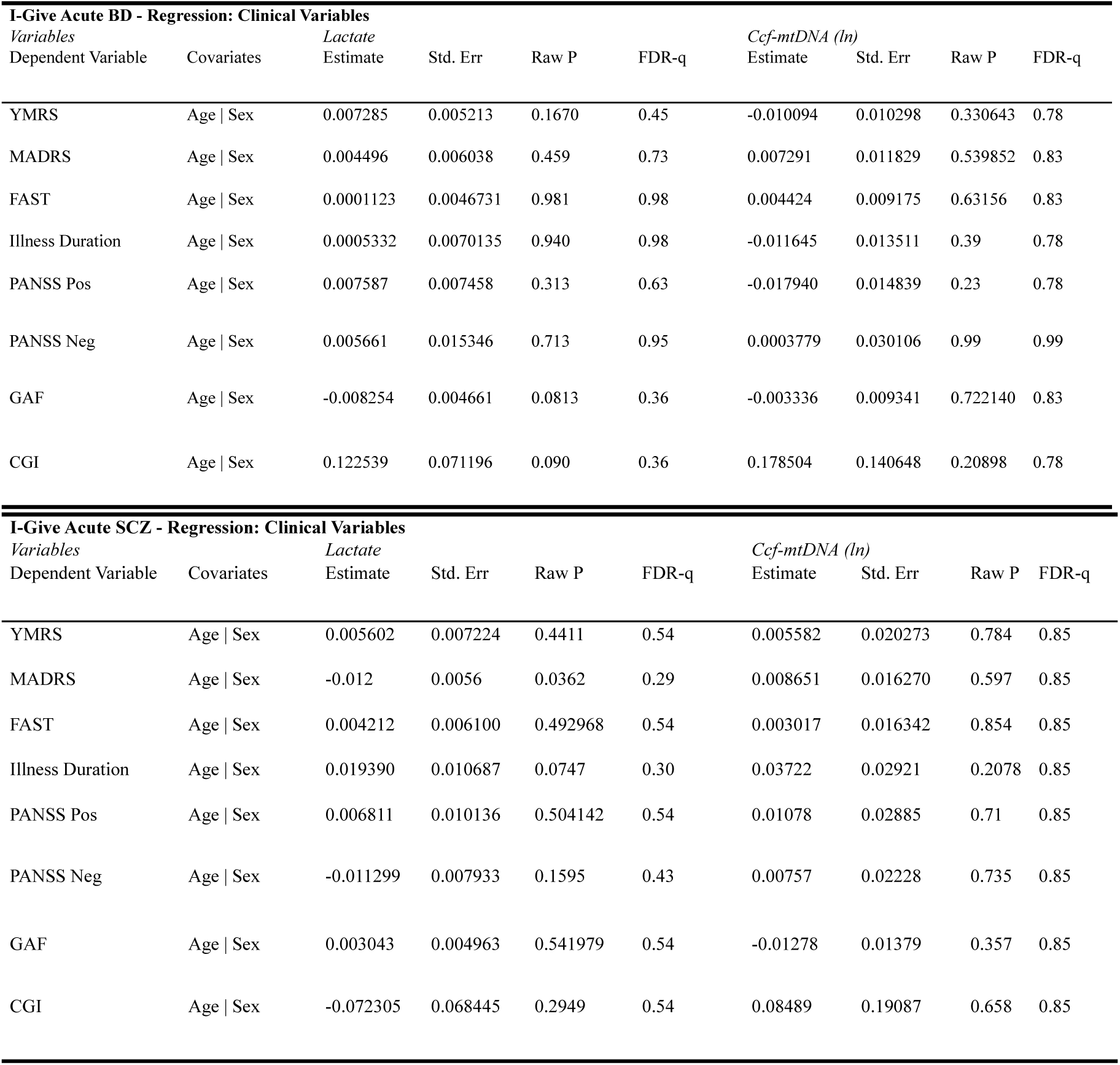
Regression analysis model for association of mitochondrial-blood based biomarkers with clinical and demographics variable in the iGIVE cohort.

**Regression analysis model for various associations of mitochondrial blood based biomarkers in the iGIVE cohort.**
| **I-Give Acute BD - Regression: Clinical Variables** |  |  |  |  |  |  |  |  |  |
| --- | --- | --- | --- | --- | --- | --- | --- | --- | --- |
| *Variables* |  | *Lactate* |  |  |  | *Ccf-mtDNA (ln)* |  |  |  |
| Dependent Variable | Covariates | Estimate | Std. Err | Raw P | FDR-q | Estimate | Std. Err | Raw P | FDR-q |
| YMRS | Age \| Sex | 0.007285 | 0.005213 | 0.1670 | 0.45 | -0.010094 | 0.010298 | 0.330643 | 0.78 |
| MADRS | Age \| Sex | 0.004496 | 0.006038 | 0.459 | 0.73 | 0.007291 | 0.011829 | 0.539852 | 0.83 |
| FAST | Age \| Sex | 0.0001123 | 0.0046731 | 0.981 | 0.98 | 0.004424 | 0.009175 | 0.63156 | 0.83 |
| Illness Duration | Age \| Sex | 0.0005332 | 0.0070135 | 0.940 | 0.98 | -0.011645 | 0.013511 | 0.39 | 0.78 |
| PANSS Pos | Age \| Sex | 0.007587 | 0.007458 | 0.313 | 0.63 | -0.017940 | 0.014839 | 0.23 | 0.78 |
| PANSS Neg | Age \| Sex | 0.005661 | 0.015346 | 0.713 | 0.95 | 0.0003779 | 0.030106 | 0.99 | 0.99 |
| GAF | Age \| Sex | -0.008254 | 0.004661 | 0.0813 | 0.36 | -0.003336 | 0.009341 | 0.722140 | 0.83 |
| CGI | Age \| Sex | 0.122539 | 0.071196 | 0.090 | 0.36 | 0.178504 | 0.140648 | 0.20898 | 0.78 |
| **I-Give Acute SCZ - Regression: Clinical Variables** |  |  |  |  |  |  |  |  |  |
| *Variables* |  | *Lactate* |  |  |  | *Ccf-mtDNA (ln)* |  |  |  |
| Dependent Variable | Covariates | Estimate | Std. Err | Raw P | FDR-q | Estimate | Std. Err | Raw P | FDR-q |
| YMRS | Age \| Sex | 0.005602 | 0.007224 | 0.4411 | 0.54 | 0.005582 | 0.020273 | 0.784 | 0.85 |
| MADRS | Age \| Sex | -0.012 | 0.0056 | 0.0362 | 0.29 | 0.008651 | 0.016270 | 0.597 | 0.85 |
| FAST | Age \| Sex | 0.004212 | 0.006100 | 0.492968 | 0.54 | 0.003017 | 0.016342 | 0.854 | 0.85 |
| Illness Duration | Age \| Sex | 0.019390 | 0.010687 | 0.0747 | 0.30 | 0.03722 | 0.02921 | 0.2078 | 0.85 |
| PANSS Pos | Age \| Sex | 0.006811 | 0.010136 | 0.504142 | 0.54 | 0.01078 | 0.02885 | 0.71 | 0.85 |
| PANSS Neg | Age \| Sex | -0.011299 | 0.007933 | 0.1595 | 0.43 | 0.00757 | 0.02228 | 0.735 | 0.85 |
| GAF | Age \| Sex | 0.003043 | 0.004963 | 0.541979 | 0.54 | -0.01278 | 0.01379 | 0.357 | 0.85 |
| CGI | Age \| Sex | -0.072305 | 0.068445 | 0.2949 | 0.54 | 0.08489 | 0.19087 | 0.658 | 0.85 |

**eTable 5.** Lactate & ccf-mtDNA across YMRS & MADRS.

**Lactate & circulating cell-free mitochondrial DNA across YMRS & MADRS.**
| *FACE-BD - BD Stable* | **YMRS**<br>**> 14**<br>**(n = 12 )** | **YMRS**<br>**8-14**<br>**(n = 44)** | **YMRS**<br>**< 8**<br>**(n = 757)** | **MADRS**<br>**< 16**<br>**(n = 175)** | **MADRS**<br>**=>16**<br>**(n = 636)** |
| --- | --- | --- | --- | --- | --- |
| Lactate | 1.14 (0.38) | 1.17 (0.46) | 1.29 (0.49) | 1.18 (0.40) | 1.18 (0.48) |
| ccf-mtDNA (ln) | 9.61 (0.71) | 9.68 (0.92) | 9.39 (0.89) | 9.54 (0.97) | 9.38 (0.87) |
| *iGIVE - BD Acute* | **YMRS**<br>**> 14**<br>**(n = 39)** | **YMRS**<br>**8-14**<br>**(n = 18 )** | **YMRS**<br>**< 8**<br>**(n = 12)** | **MADRS**<br>**< 16**<br>**(n = 30)** | **MADRS**<br>**=>16**<br>**(n = 39)** |
| Lactate | 1.53 (0.43) | 1.44 (0.64) | 1.46 (0.48) | 1.48 (0.57) | 1.50 (0.43) |
| ccf-mtDNA (ln) | 9.84 (1.16) | 10.2 (0.93) | 9.82 (0.97) | 10.0 (0.84) | 9.86 (1.20) |
| *iGIVE - SCZ Acute* | **YMRS**<br>**> 14**<br>**(n = 39)** | **YMRS**<br>**8-14**<br>**(n = 18 )** | **YMRS**<br>**< 8**<br>**(n = 12)** | **MADRS**<br>**< 16**<br>**(n = 30)** | **MADRS**<br>**=>16**<br>**(n = 39)** |
| Lactate | 1.51 (0.43) | 1.59 (0.29) | 1.49 (0.56) | 1.34 (0.42) | 1.60 (0.46) |
| ccf-mtDNA (ln) | 9.63 (1.26) | 9.58 (1.37) | 9.72 (1.23) | 9.56 (1.24) | 9.70 (1.27) |

### Supplementary Figures

**eFigure 1:**
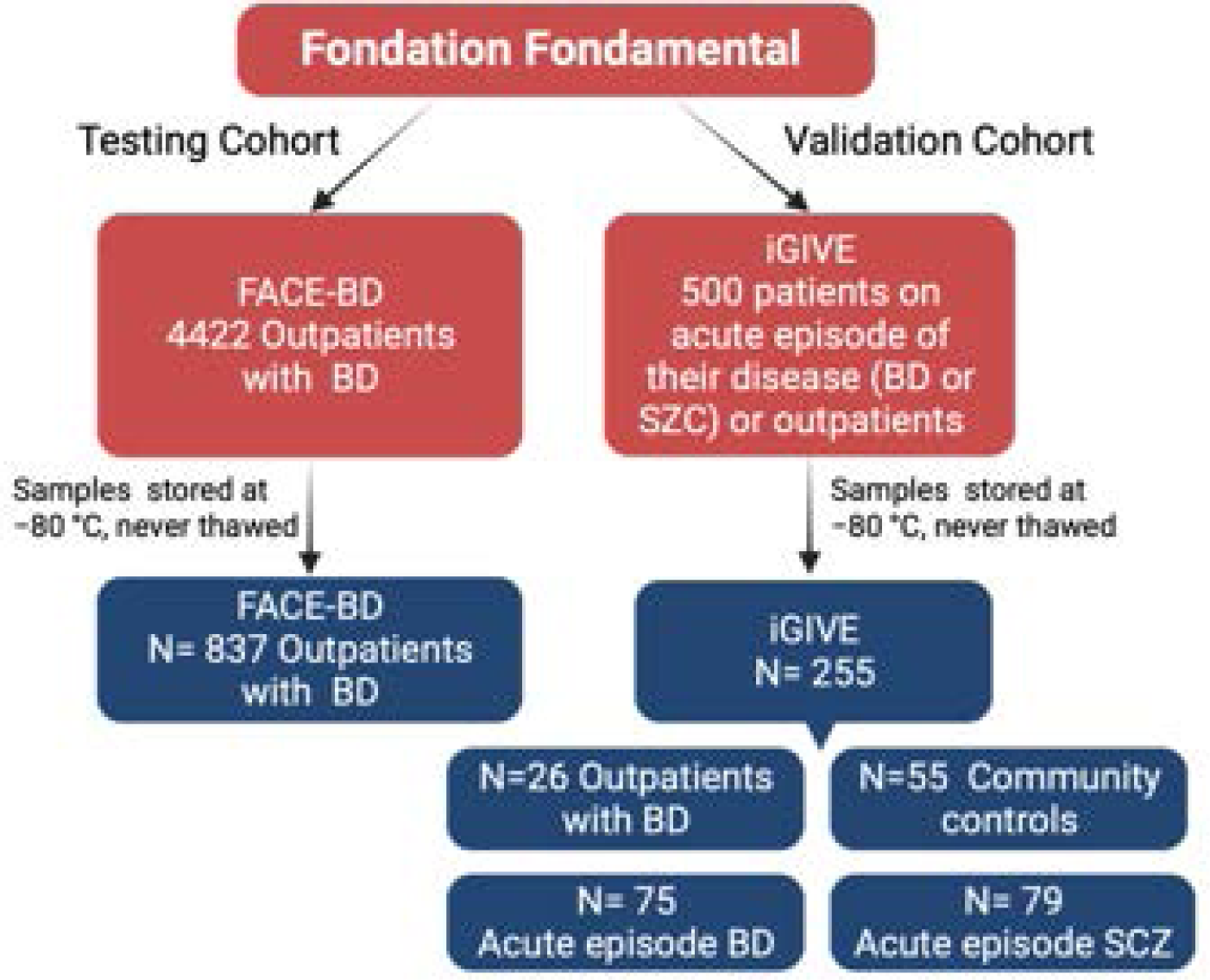
Testing cohort (FACE-BD) and validation cohort (i-GIVE) sample inclusion.

**eFigure 2:**
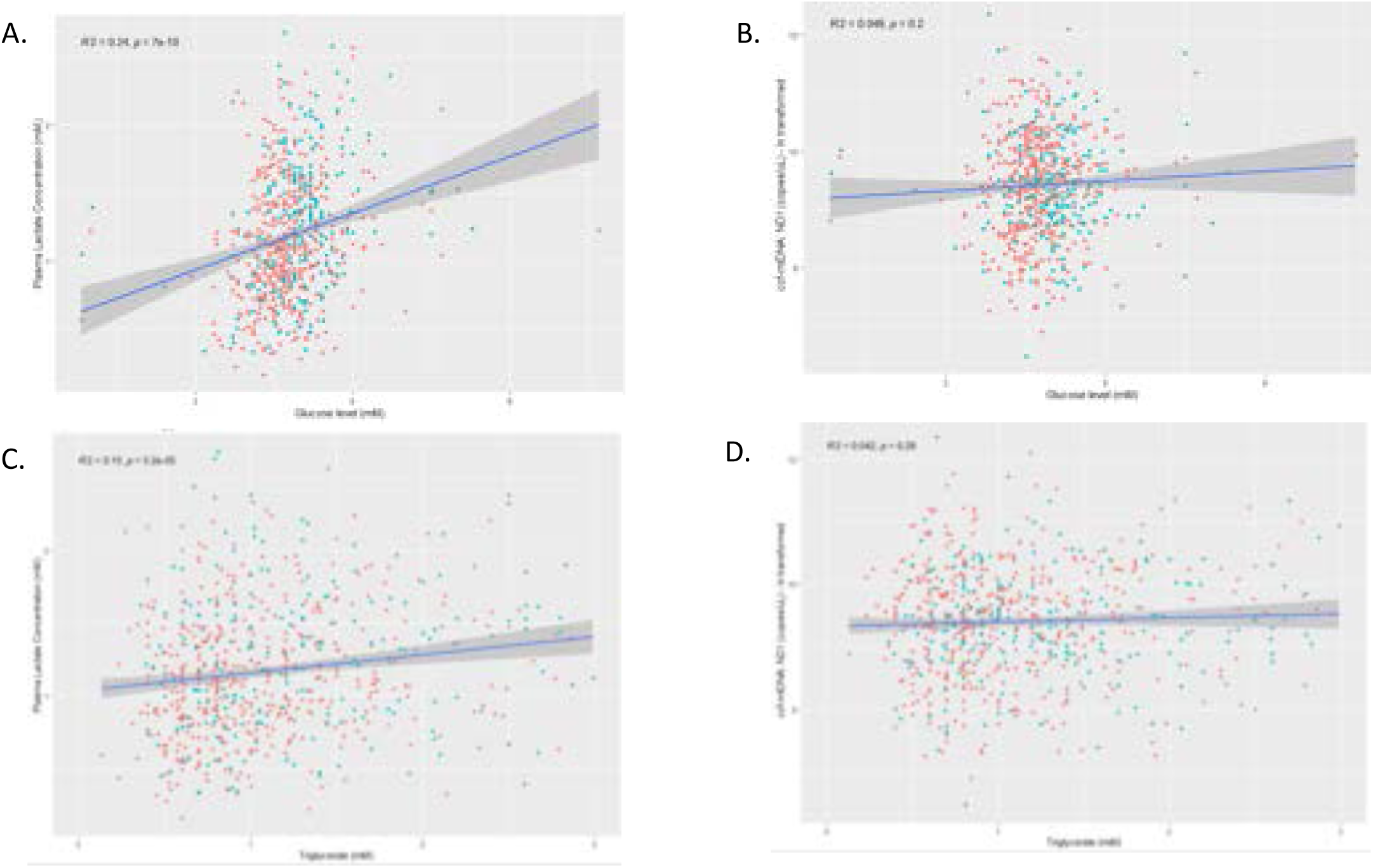
Association of peripheral levels of lactate (A, C) and circulating cell free mitochondrial DNA (B, D) with fasting glucose levels and triglycerides in stable patients with bipolar disorder.

**eFigure 3.**
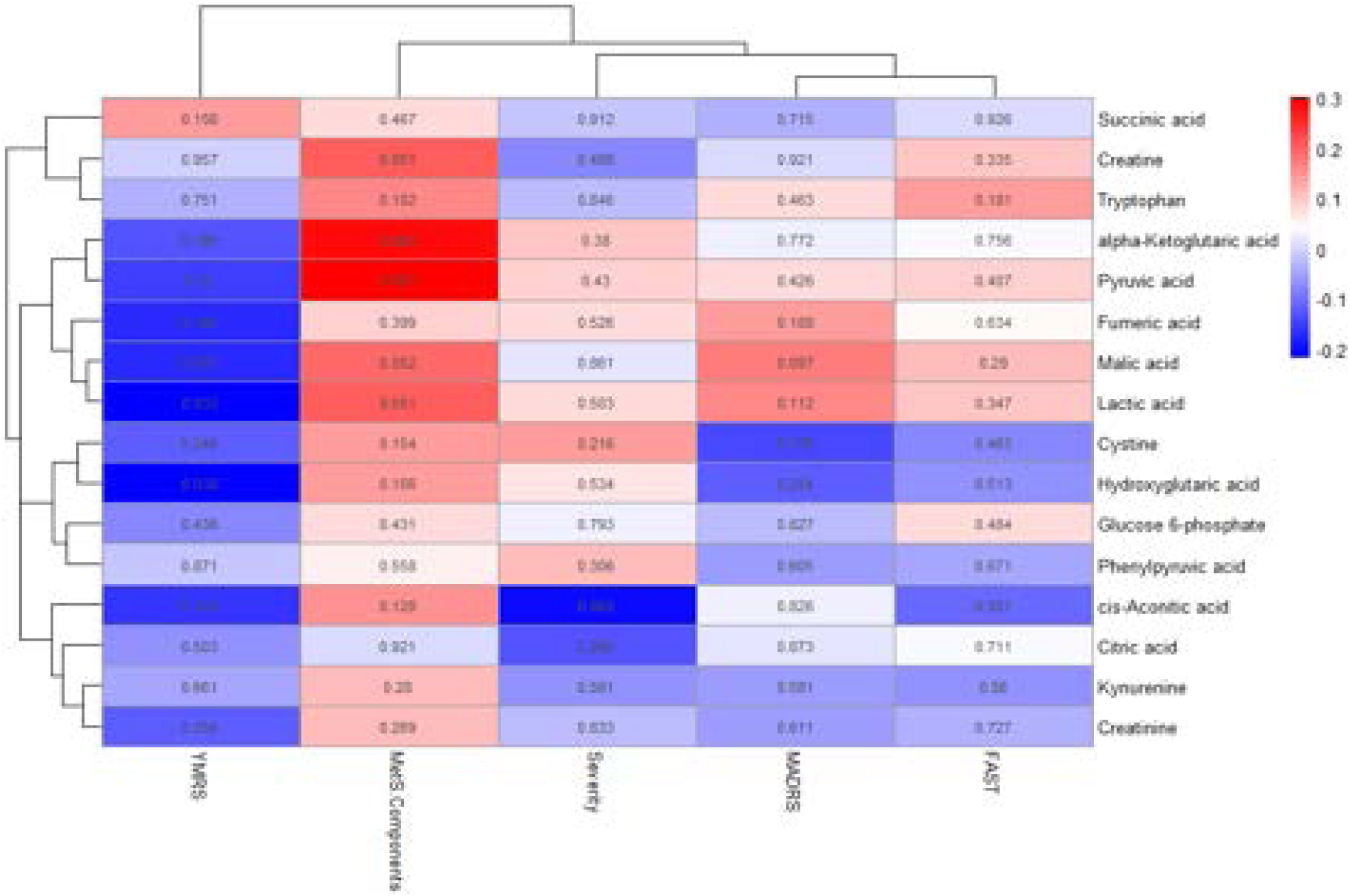
Clustering analysis of metabolites and clinical scales.

**eFigure 4.**
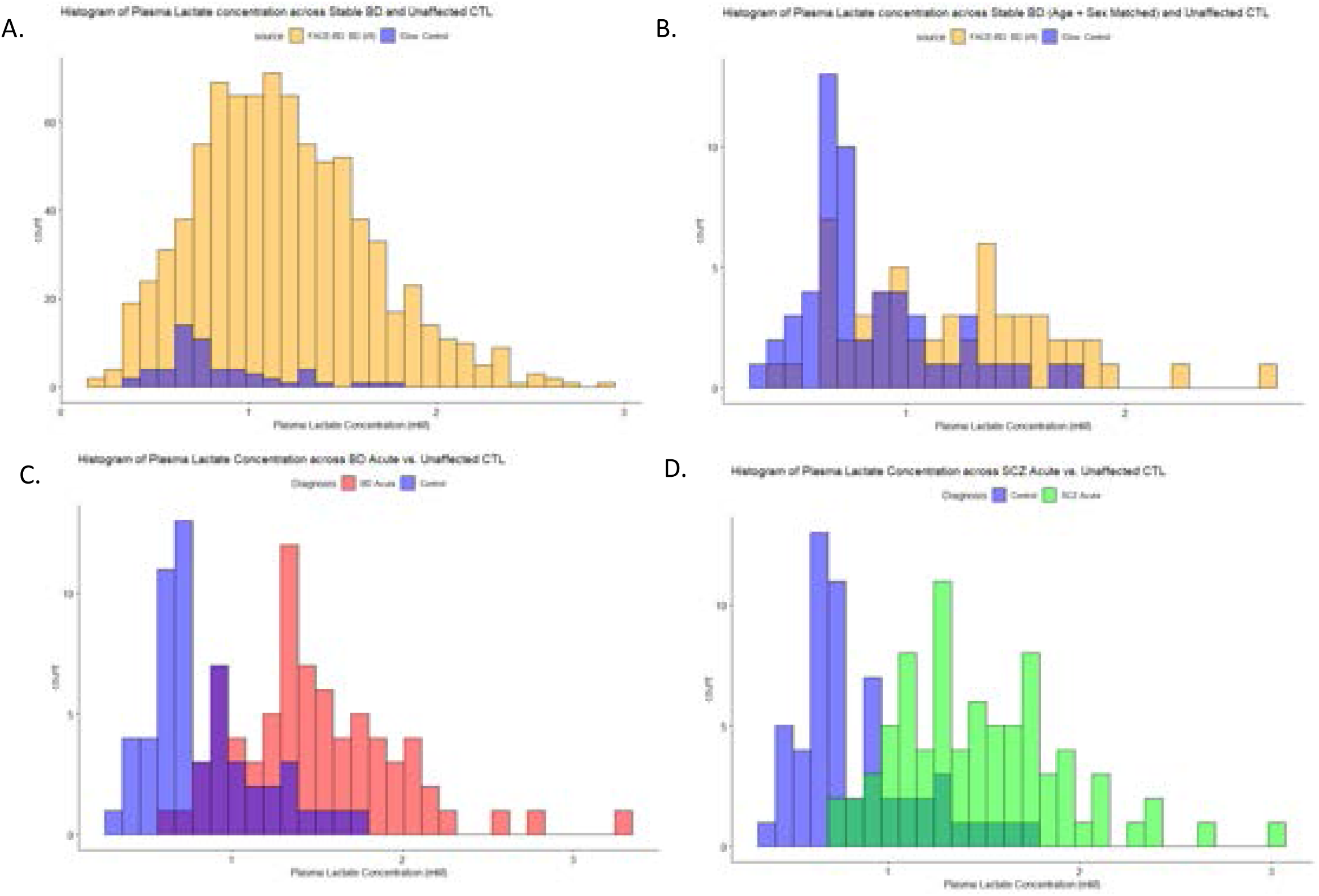
Distribution of plasma lactate concentration across non-psychiatry controls, stable BD age and sex matched (A, B), acute BD (C) and acute SCZ (D).

### Supplemental Methods

#### Extraction and detection of circulating-cell free mitochondrial DNA

The extraction of ccf-mtDNA from plasma was conducted utilizing the QIAamp DNA Mini kit based on the manufacturer’s protocol for DNA Purification from Blood or Bodily fluids using spin columns. To collect ccf-mtDNA, we used 50μL of plasma. To elute from the column, we used 100 μL of UltraPure distilled water, free of DNAase and RNAase (Invitrogen). HEK DNA was previously extracted and quantified using digital PCR to allow us to determine the absolute concertation (copies/uL) of ccf-mtDNA in patient plasma samples. The HEK DNA was of known concentration, and was diluted in half-logs, of up to twelve standard points. To detect ccf-mtDNA and represent both the major and minor arc of the mitochondrial genome, ND4 and ND1 were used respectively, with PPIA and B2M as respective nuclear controls. The qPCRs were run as a duplex reaction using a 20μL mixture made up of 10μL TaqMan Fast Advanced Master Mix, 4μL DNA and 1μL of each forward primers, reverse primers and TaqMan probe for each gene. All qPCRs were run using BioRad’s C1000 Thermal cycle CFX384 Real Time System following cycling conditions described by TaqMan manufacturer: 50°C for 2 minutes, 95°C for 20 seconds, 40 cycles of 95°C for 3 seconds and 60°C for 30 seconds, followed by a fluorescent read per cycle. Absolute concentrations are determined against the 12-point standard curve using linear equation. was assayed in technical triplicate and concentrations were calculated using a standard curve. Experimenters were blinded when measuring ccf-mtDNA and conducting raw analyses. A strong correlation between levels of ND1 and ND4 is show below, highlining the reliability of ccf-mtDNA in detecting mitochondrial circulating DNA.

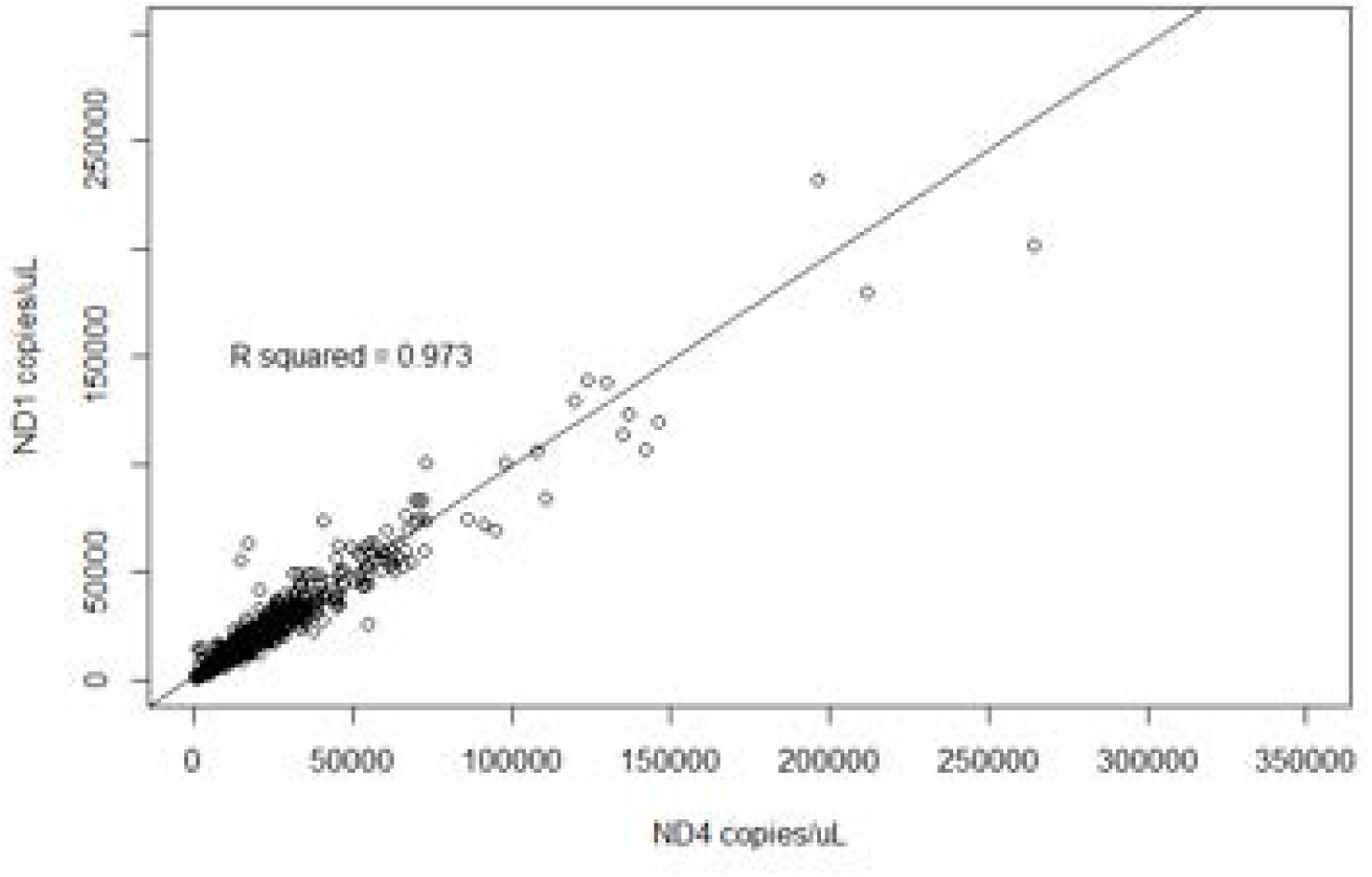

